# Improving prediction of compound function from chemical structure using chemical-genetic networks

**DOI:** 10.1101/112698

**Authors:** Hamid Safizadeh, Scott W. Simpkins, Justin Nelson, Chad L. Myers

## Abstract

The drug discovery process can be significantly improved through understanding how the structure of chemical compounds relates to their function. A common paradigm that has been used to filter and prioritize compounds is ligand-based virtual screening, where large libraries of compounds are queried for high structural similarity to a target molecule, with the assumption that structural similarity is predictive of similar biological activity. Although the chemical informatics community has already proposed a wide range of structure descriptors and similarity coefficients, a major challenge has been the lack of systematic and unbiased benchmarks for biological activity that covers a broad range of targets to definitively assess the performance of the alternative approaches.

We leveraged a large set of chemical-genetic interaction data from the yeast *Saccharomyces cerevisiae* that our labs have recently generated, covering more than 13,000 compounds from the RIKEN NPDepo and several NCI, NIH, and GlaxoSmithKline (GSK) compound collections. Supportive of the idea that chemical-genetic interaction data provide an unbiased proxy for biological functions, we found that many commonly used structural similarity measures were able to predict the compounds that exhibited similar chemical-genetic interaction profiles, although these measures did exhibit significant differences in performance. Using the chemical-genetic interaction profiles as a basis for our evaluation, we performed a systematic benchmarking of 10 different structure descriptors, each combined with 12 different similarity coefficients. We found that the All-Shortest Path (ASP) structure descriptor paired with the Braun-Blanquet similarity coefficient provided superior performance that was robust across several different compound collections.

We further describe a machine learning approach that improves the ability of the ASP metric to capture biological activity. We used the ASP fingerprints as input for several supervised machine learning models and the chemical-genetic interaction profiles as the standard for learning. We found that the predictive power of the ASP fingerprints (as well as several other descriptors) could be substantially improved by using support vector machines. For example, on held-out data, we measured a 5-fold improvement in the recall of biologically similar compounds at a precision of 50% based upon the ASP fingerprints. Our results generally suggest that using high-dimensional chemical-genetic data as a basis for refining chemical structure descriptors can be a powerful approach to improving prediction of biological function from structure.

## INTRODUCTION

Discovery, design, and development of new drugs that reveal desired and reproducible biochemical behavior against a particular biomolecular target with minimal side effects are challenging. Despite the scientific and technological advances in drug discovery during the past 60 years, the number of drugs approved per billion US dollars that were spent for the development of novel drugs has halved roughly every 9 years since 1950 (Eroom’s Law in contrast to Moore’s Law) [1]. Following the similar property principle (SPP) [2], Ligand-based virtual screening (LBVS) has been commonly used as an a priori step to high-throughput screening (HTS) [3,4,5] to rank compounds of a large database in the decreasing order of their similarity to a reference or target molecule with known biological activity (**Fig. 1**). According to the similar property principle, structurally similar molecules are more likely to represent similar biological activities and physicochemical properties. Although there are limitations to the similar property principle [6], such as the case of activity cliffs where a very small modification in the structure of a molecule may drastically alter its biological properties [7], this structure-activity relationship is broadly consistent throughout the larger flat regions of activity landscapes [8,9]. Hence, the need for high performance structural similarity that extracts structurally analogous compounds from a database is inevitable.

**Figure 1.**
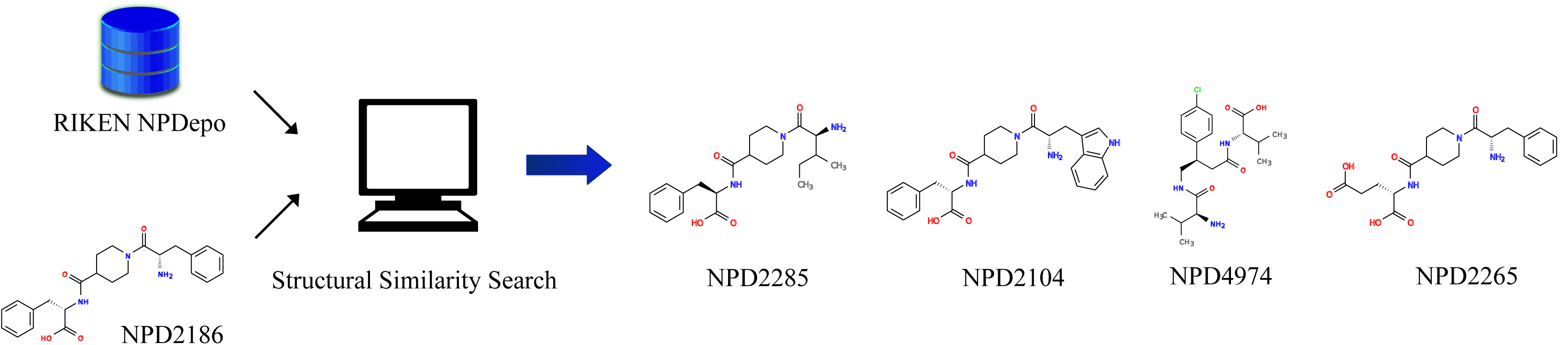
Ligand-based virtual screening of a target (e.g., NPD2186 from RIKEN NPDepo). All the compounds of the MOSAIC database (http://mosaic.cs.umn.edu) [29] were ranked in the decreasing order of structural similarity to the target molecule based upon the similar property principle (SPP). In this ranked list, NPD4974 had a very distinct chemical-genetic profile from NPD2186, appearing as a false positive generated by SPP. Here, we described all the compounds using ASP fingerprints (depth 8) and measured the structural similarity using the Braun-Blanquet similarity coefficient.

To accelerate the retrieval of the compounds of a desired class that are active against a protein target, the chemical informatics community has suggested a wide range of structure descriptors and similarity coefficients that are able to extract candidates with similar biological activity from structural compound libraries. The most widely used representation of molecular graphs in these expanding databases of two- and three-dimensional molecular structures is based upon chemical fingerprints [10,11,12], where a molecular graph is represented by a fixed-length bit-vector that enumerates all the bounded-length paths in the graph and encodes the presence or absence of substructural fragments. The degree of similarity of two structural vectors describing two different compounds is usually measured by similarity coefficients, among which the well-known Tanimoto coefficient has still remained the coefficient of choice to capture the highest level of intermolecular similarity and thus biological activity [11,13]. The Tanimoto coefficient, which is formulated as the number of features shared between two molecules divided by the total number of features presented in both molecules, offers a suitable degree of chemical similarity between compounds, although this coefficient suffers from an intrinsic bias towards selection of smaller compounds [14,15]. However, the research community has lacked a systematic benchmark that assesses the performance of these structure descriptors and similarity coefficients over a broad range of protein targets in an unbiased manner.

Chemical genomic approaches, which focus on the systematic mapping of chemical-genetic interactions, offer a valuable new source of data to connect structure to function. These chemical-genetic maps take advantage of the massive wealth of chemical-genetic interaction profiles. The yeast Saccharomyces cerevisiae is a well-characterized eukaryotic system for which the genome-wide gene deletion project has identified ∼5000 viable deletion mutants [16]. Testing each one of these viable mutants for hypersensitivity to a bioactive compound generates a chemical-genetic interaction profile in which the relative fitness of a selected group of mutant strains with defined genetic perturbations in response to the bioactive compound is quantified [17,18]. These chemical-genetic interaction profiles provide functional information for a compound that can be interpreted through the global genetic interaction network mapped for the yeast [19]. If a bioactive compound inhibits a target protein, the loss-of-function mutations in a gene that encodes the protein models the primary effects of the compound, and the genetic interaction profile of the target gene resembles the chemical-genetic interaction profile of the compound that inhibits the target pathway. Consequently, the chemical-genetic interaction profiles of bioactive compounds can link those compounds to their cellular target pathways in an unbiased manner. These profiles can be annotated to specific biological processes to predict the general mechanisms of action for the bioactive compounds and can serve as an unbiased genome-wide measure for their biological activity [20].

We generated a systematic benchmark based upon the similar property principle to assess the performance of several structure descriptors and similarity coefficients in prediction of the chemical-genetic interaction profiles of hundreds of compounds, with the assumption that these profiles provided an unbiased genome-wide measure of biological activity. We generated and annotated the yeast chemical-genetic interaction profiles for more than 13,000 compounds from the RIKEN NPDepo as well as several NCI/NIH/GSK compound collections [20]. We systematically benchmarked 10 different structure descriptors, each combined with 12 different similarity coefficients, to identify the pair with the superior prediction of biological activity by using the chemical-genetic interaction profiles as a basis for the biological activity of our compounds. We further developed several supervised machine learning models to improve our prediction of the biological activity of compounds from chemical structures, gaining higher predictive power that was not in the ability of similarity coefficients. We found that support vector machines (SVMs) [21] can significantly enhance the power of our chemical fingerprints for predicting the biological activity of compounds.

## RESULTS AND DISCUSSION

To evaluate the performance of the commonly used structure descriptors and similarity coefficients in predicting the biological activity of compounds, we exhaustively searched for all the pairwise candidates (i.e., one structure descriptor and one similarity coefficient) that provided high predictive power over a wide range of protein targets using our chemical-genetic interaction data. We generated, in our labs, the chemical-genetic interaction profiles for 13,524 compounds from several diverse compound collections [20]. We used a subset of these screened compounds that exhibited high confidence predictions for the annotated processes and biological pathways based upon our chemical-genetic interaction profiles [20,22]. We included in our benchmarking system two compound sets independently: (1) 826 compounds from the RIKEN Natural Product Depository (NPDepo) compound collection, which we called RIKEN high confidence collection. (2) 659 compounds from several NCI/NIH/GSK collections, which we called the NCI/NIH/GSK high confidence collection (see **Material**).

### Establishing a systematic benchmark for chemical similarity measures

We aimed at predicting the compound pairs that exhibited the most similar function based upon our chemical-genetic interaction profiles, which served as an unbiased genome-wide measure of biological activity. We labeled only 10% of the compound pairs with the highest (cosine) profile similarity as our gold standard for true positives, which were highly prioritized in our systematic benchmark. Following the similar property principle, a large number of these true positives should be identified from the structural similarity of our compounds.

Our benchmarking system consisted of two main components: structure descriptors and similarity coefficients. We evaluated both components through our systematic benchmark to find the best performer of each component for prediction of the biological activity of our compounds. We used jCompoundMapper [23] to describe all our compounds in 10 different structure spaces (**Table 1**), where a compound was described with a fixed-length bit-vector that indicated the presence or absence of a certain number of fingerprints. The number of features required for the description of the compounds in a structure space varied based upon the space definition and collection properties (e.g., only 9 of the predefined features in the MACCS keys were found in the RIKEN high confidence collection, while RAD2D fingerprints generated 91082 features to describe this compound collection). We used 12 widely used similarity coefficients (**Table 2**) to measure the degree of similarity of two compounds described by a given structure descriptor.

**Table 1.** Structure descriptors. 10 different topological, fingerprint-based structure descriptors generated by jCompoundMapper for the description of each compound in our datasets. The right column represented the total number of features needed to describe our high confidence RIKEN collection (826 compounds).

| **ID** | **Name** | **Description** | **# Features** |
| --- | --- | --- | --- |
| SD1 | AP2D | Topological atom pairs | 1210 |
| SD2 | ASP | All-shortest path encodings | 26114 |
| SD3 | AT2D | Topological atom triplets | 56900 |
| SD4 | DFS | All-path encodings | 48267 |
| SD5 | ECFP | Extended connectivity fingerprints | 42131 |
| SD6 | LSTAR | Local path environments | 84450 |
| SD7 | MACCS | Predefined pharmacophores | 9 |
| SD8 | PHAP2POINT2D | Topological pharmacophore pair encodings | 17 |
| SD9 | PHAP3POINT2D | Topological pharmacophore triplet encodings | 302 |
| SD10 | RAD2D | Topological molprint-like fingerprints | 91082 |

**Table 2.** Similarity coefficients. 12 different similarity coefficients (several of these coefficients were collected by Raymond and Willett [12]) for measurement of the degree of similarity of two compounds described by a given fingerprint-based structure descriptor. Here, *x* = number of bits set in both fingerprints, *y* = number of bits set in the first fingerprint, *z* = number of bits set in the second fingerprint, and *w =* number of bits in the bit string. For the Tullos similarity coefficient, 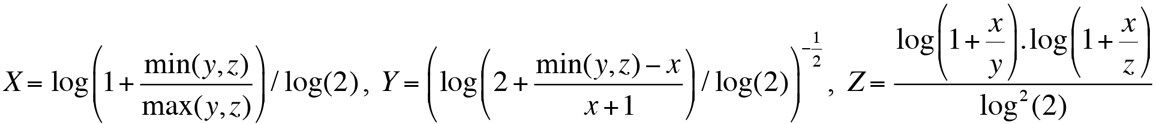

| ID | Name | Measurement | Range |
| --- | --- | --- | --- |
| SC1 | Cosine | $\frac{x}{\sqrt{yz}}$ | 0 to 1 |
| SC2 | Tanimoto | $\frac{x}{y+z-x}$ | 0 to 1 |
| SC3 | Kulczynski | $\frac{x(y+z)}{2yz}$ | 0 to 1 |
| SC4 | Dice | $\frac{2x}{y+z}$ | 0 to 1 |
| SC5 | Sokal/Sneath | $\frac{x}{2y+2z-3x}$ | 0 to 1 |
| SC6 | Tullos | $XYZ$ | 0 to 1 |
| SC7 | McConnaughey | $\frac{x(y+z)-yz}{yz}$ | -1 to 1 |
| SC8 | Asymmetric | $\frac{x}{\min(y,z)}$ | 0 to 1 |
| SC9 | Braun-Blanquet | $\frac{x}{\max(y,z)}$ | 0 to 1 |
| SC10 | Russel/Rao | $\frac{x}{w}$ | 0 to 1 |
| SC11 | Euclidean | $\frac{1}{1+\sqrt{y+z-2x}}$ | 0 to 1 |
| SC12 | Dot-product | $x$ | 0 to ∞ |

### Evaluating the performance of chemical similarity measures

We exhaustively searched for the best-performing chemical similarity measure (i.e., one structure descriptor and one similarity coefficient) in predicting the biological activity of our compounds. We described all our compounds in 10 different structure spaces and measured the structural similarity of two compounds described in a given structure space by using a coefficient of structural similarity, generating compound similarities for all the combinations of structure descriptors and similarity coefficients. We ranked all these scores of structural similarity for the prediction of the chemical-genetic profile similarity of our compounds and measured precision at many recalls to evaluate the performance of alternative models (**Suppl. Table 1**). To isolate the winning structure descriptor, we looked at the precision of all the prediction models at several (lower) recalls for our RIKEN high confidence collection, which distinguished ASP, LSTAR, and RAD2D fingerprints as the structure descriptors with superior predictive power (**Fig. 2a**). The wide range of precision values achieved by different structure descriptors, assuming that a single similarity coefficient was used, showed that our chemical-genetic interaction profiles could highly separate structure descriptors in terms of their efficiency for the prediction of the biological activity of compounds. We compared the relative performance of our distinguished models with the predictive power of extended-connectivity fingerprints (ECFP) [24], which has recently been one of the most common descriptors to represent molecular graphs. However, ECFP did not generally outperform ASP, LSTAR, and RAD2D fingerprints, except at very few recalls (**Fig. 2b**). Because this finding might simply be a result of our collection property, we used our NCI/NIH/GSK high confidence collection to validate the predictive performance of the ASP, LSTAR, and RAD2D fingerprints, which strongly confirmed the superiority of these descriptors over ECFP at many recalls (**Fig. 3**).

**Figure 2.**
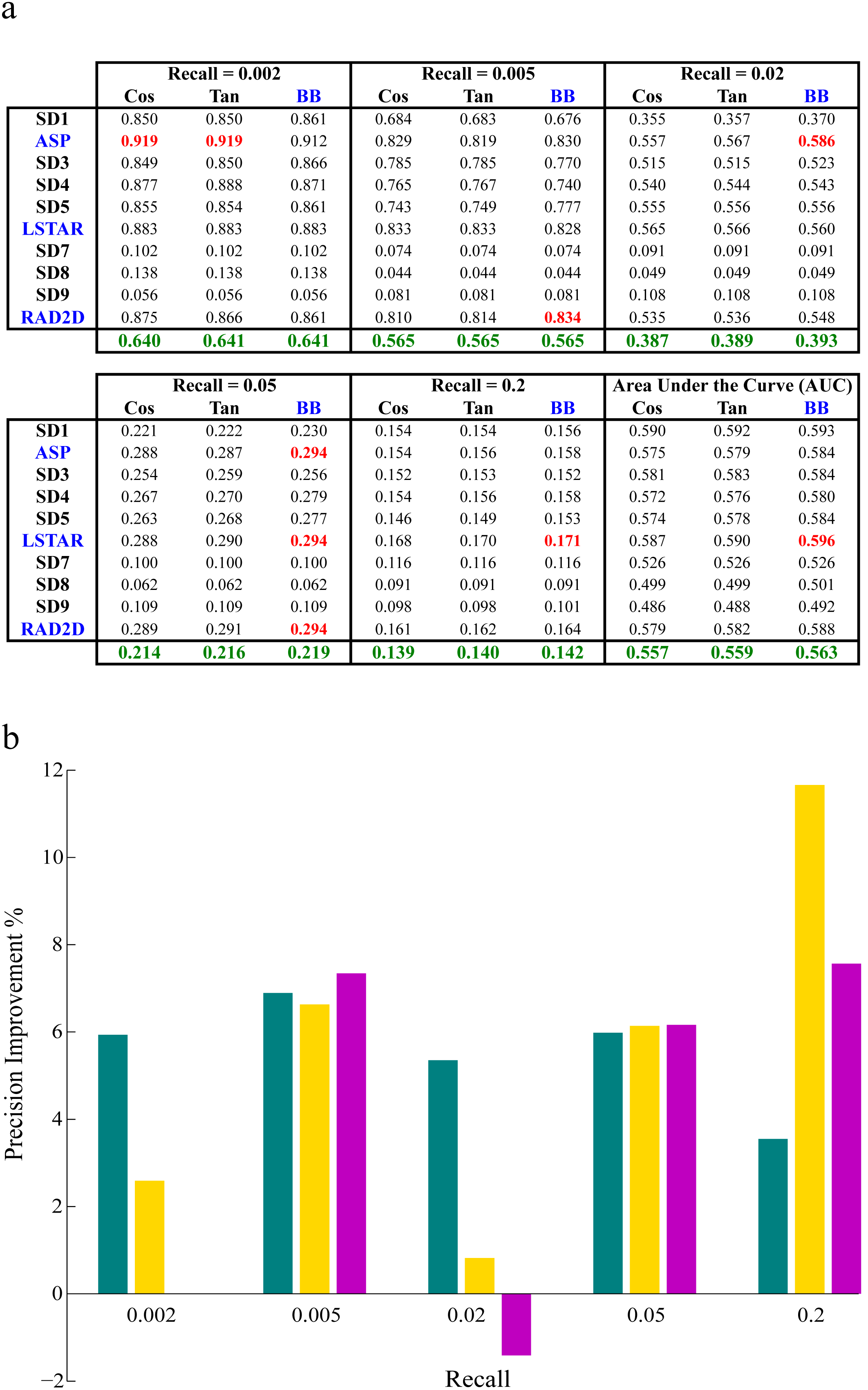
Model precision for all structure descriptors paired with the Cosine, Tanimoto, or Braun-Blanquet similarity coefficient using our RIKEN high confidence data collection. **(a)**Precision at several recalls and the area under the ROC curve for each model. The red and green values represented the highest precision achieved at a given recall and the average precision over all the structure descriptors for a given similarity coefficient at a recall, respectively. **(b)** Relative performance of ASP (teal), LSTAR (gold), and RAD2D (magenta) fingerprints to ECFP. For all the structure descriptors that required a depth of description, precision was measured at depth 8.

**Figure 3.**
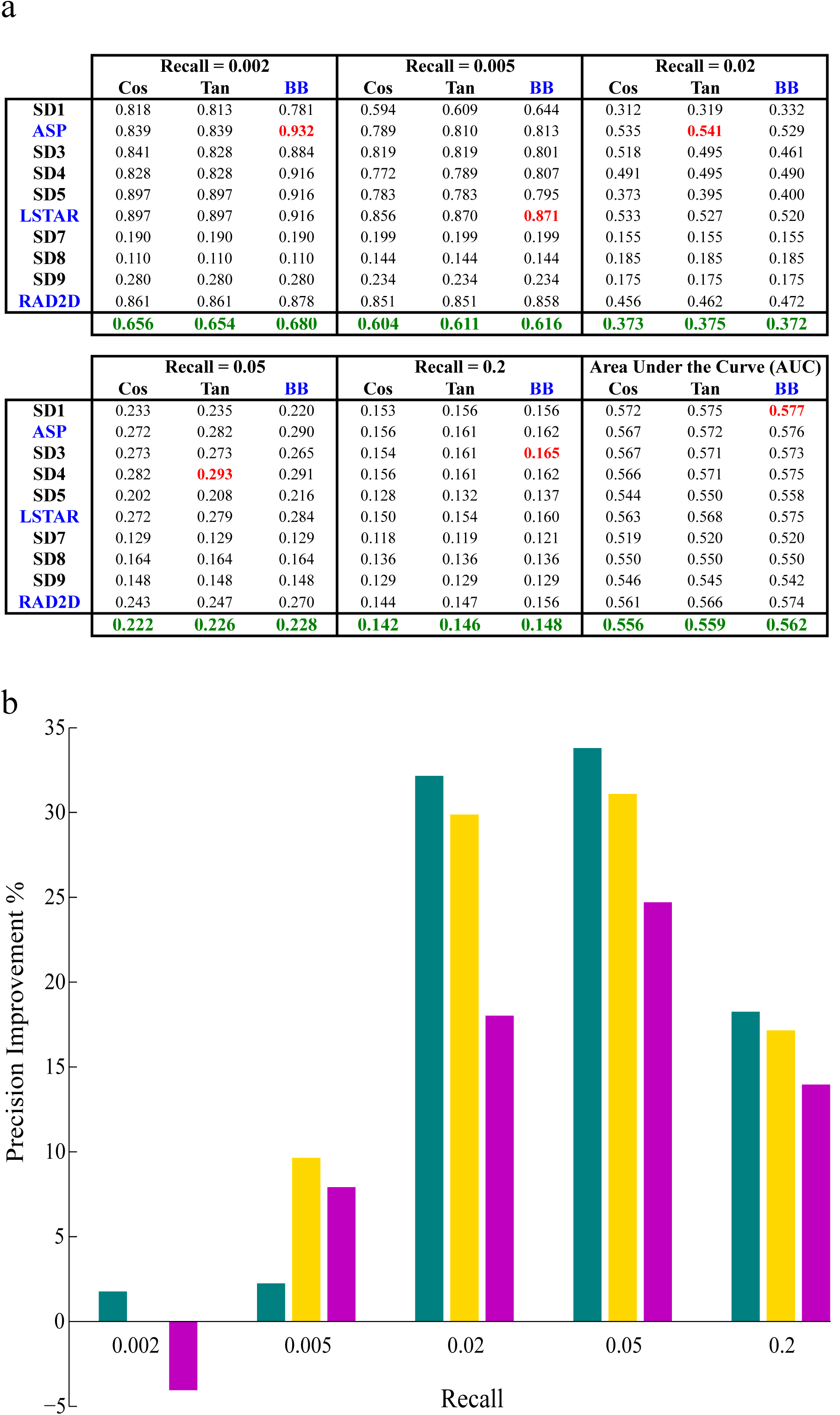
Model precision for all structure descriptors paired with the Cosine, Tanimoto, or Braun-Blanquet similarity coefficient using our NCI/NIH/GSK high confidence data collection. **(a)** Precision at several recalls and the area under the ROC curve for each model. The red and green values represented the highest precision achieved at a given recall and the average precision over all the structure descriptors for a given similarity coefficient at a recall, respectively. **(b)** Relative performance of ASP (teal), LSTAR (gold), and RAD2D (magenta) fingerprints to ECFP. For all the structure descriptors that required a depth of description, precision was measured at depth 8.

ASP fingerprints encoded a graph traversal over all atoms in a molecular graph but stored only the shortest paths between atoms, whereas LSTAR and RAD2D fingerprints described the radial environment of all atoms in the molecular graph [23]. As a result, the ASP encoding that described our compound collections needed fewer features than LSTAR and RAD2D encodings (**Table 1**) although the predictive performance of the ASP fingerprints was higher or comparable with that of LSTAR and RAD2D fingerprints at several recalls. Moreover, LSTAR fingerprints generally exhibited higher performance than RAD2D fingerprints at several recalls (**Figs. 2–3**), which could be justified by additional information that LSTAR fingerprints collected from the radial environment of the atoms by definition. Therefore, we determined ASP and LSTAR fingerprints as the winning structure descriptors to describe compounds and predict the ones with the highest biological similarity to a target molecule using a similarity coefficient.

We systematically benchmarked 12 similarity coefficients (**Table 2**) by using our RIKEN high confidence collection and measured the predictive performance of every coefficient over all structure descriptors to find the winning similarity coefficient. We found that several (8 out of 12) coefficients were able to exhibit consistent high performance across all structure descriptors, although 4 similarity coefficients (Asymmetric, Russel/Rao, Euclidean, and Dot-product) failed in some structure descriptor spaces because their precision significantly dropped at lower recalls. In other words, precision of several models (each corresponding to one structure descriptor) was substantially low for each of these 4 similarity coefficients at many lower recalls (**Suppl. Table 1**), which indicated that these 4 coefficients were unable to predict the biological similarity of our compounds across different structure descriptors consistently. We, as a result, removed these 4 coefficients from our analysis and focused only on 8 remaining similarity coefficients. We found that the Braun-Blanquet similarity coefficient [25] resulted in the higher precision at many recalls compared to all other coefficients, including Tanimoto and cosine coefficients (**Fig. 2a**), which have been widely used by the chemical informatics community. For the Braun-Blanquet similarity coefficient, the average precision and the average area under the receiver operating characteristic (ROC) curves across all structure descriptors were slightly higher at many recalls compared to those of the Tanimoto and cosine coefficients (**Fig. 2a**; columnar green values), suggesting that this simple coefficient of structural similarity could confidently be used in place of the traditional Tanimoto coefficient for the ranking of database compounds in the decreasing order of biological similarity to a target molecule. The Braun-Blanquet coefficient, which was simply formulated as the number of features common between two molecules divided by the total number of features presented by the larger molecule, determined the degree of contribution of the smaller molecular graph to the larger one. We further measured the performance of our predictive models using our NCI/NIH/GSK high confidence collection to validate the superiority of the Braun-Blanquet coefficient over other similarity coefficients (**Fig. 3a** and **Suppl. Table 2**). We, therefore, paired the Braun-Blanquet similarity coefficient with the ASP and LSTAR structure descriptors as our predictive models for ligand-based virtual screening.

### Optimizing the depth of structure descriptors

One major parameter involved in the structural description of compound collections was the describing depth; a high depth generated numerous features to cover the global environment of each atom, whereas a low depth only focused on describing the local neighborhood of atoms in the molecular graph. We assessed the impact of the depth of 5 structure descriptors (ASP, DFS, ECFP, LSTAR, and RAD2D) in predicting the biological activity of our compounds using the Braun-Blanquet similarity coefficient (**Fig. 4**). Describing our RIEKN high confidence collection at a high depth generally resulted in strong predictions at lower recalls but moderate outcome at higher recalls because a high depth was able to predict the compound pairs that were structurally and therefore functionally very similar according to the similar property principle (SPP) but also pushed undesired pairs such as activity cliffs to the top of our predictions. On the other hand, a low describing depth was able to capture the similarity of two compounds in the local chunks of the two molecular graphs that were essential for functional similarity, which eventually resulted in drawing reasonable predictions at lower recalls. We, furthermore, evaluated these results using our NCI/NIH/GSK high confidence collection (**Suppl. Fig. 1**), which confirmed similar general trends that impacted our predictions at several recalls using 10 different describing depths. Therefore, the structural description of a compound collection at high depths not only was unnecessary and inefficient but also increased the computation time and space complexity. We selected depth 8 for our evaluations although other neighboring depths were also justifiable.

**Figure 4.**
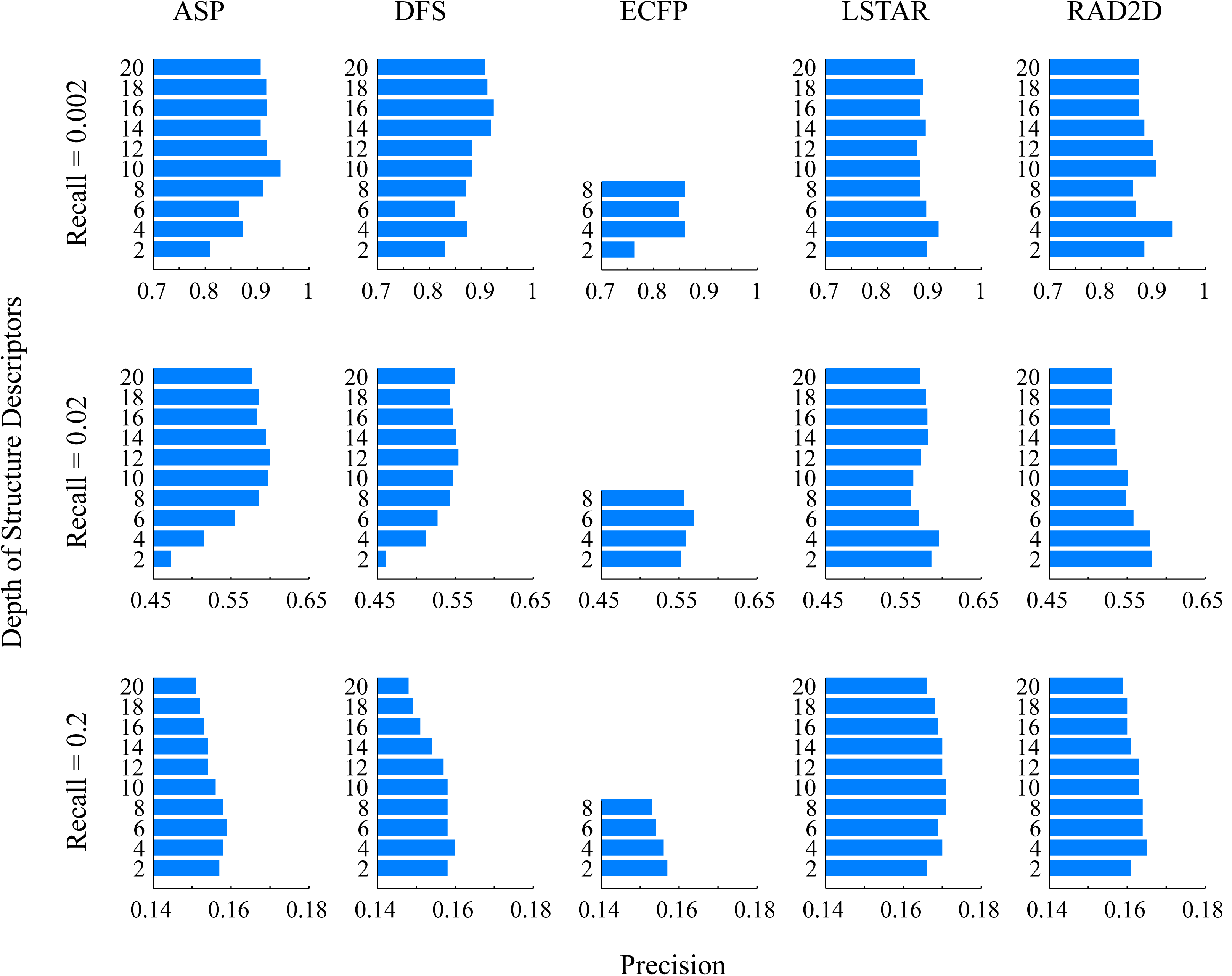
Depth impact of structure descriptors on the performance of our prediction models. We measured the precision of our prediction models at 10 consequent molecular depths for 5 different structure descriptors, each paired with the Braun-Blanquet similarity coefficient, using our RIKEN high confidence collection.

### Improving the prediction performance via SVM models

To increase the ability of chemical structures in predicting the biological activity of our compounds, we designed several supervised machine learning models and took advantage of the great wealth of chemical-genetic interaction maps for supervision. Moreover, we used supervised principal component analysis [26] via chemical-genetic interaction maps to extract a number of features from the more informative compound substructures that highly related structural data to the biological activity of our compounds. We found that support vector regression (SVR) models [21] were able to boost the prediction performance of the functional activity of our compounds from their chemical structures by weighting supervised principal components, where chemical-genetic interaction profiles were also input to the learning models for supervision.

We designed a learning pipeline (**Fig. 5a**) to predict chemical-genetic profile similarities by creating bootstraps [27] and pairwise structural encodings (see **Methods**). We implemented support vector regression models in LibSVM [28], a popular open source support vector machine learning library developed at National Taiwan University, and used Radial Basis Function (RBF) kernels for building our epsilon support vector regression models. We developed precision-recall (PR) curves to evaluate the performance of our models for different structure descriptors, where only 10% of the compound pairs from our RIKEN high confidence collection with the highest chemical-genetic profile similarity were labeled as the gold standard for true positives. We found that a subset of our structure descriptors (SD1-SD6 and SD10) were able to achieve significantly higher performance in the prediction of the functional similarity of our compounds than the best-performing chemical similarity measures (i.e., ASP or LSTAR fingerprints along with the Braun-Blanquet similarity coefficient). The learning curves for the ASP and LSTAR fingerprints (**Fig. 5b**) exhibited that we were able to gain a 5-fold improvement in the recall of biologically similar compounds at a precision of 50%. However, the degree of improvement was dependent on the functional diversity of datasets, which could result in modestly higher performance for particular collections with higher diversity; for instance, we improved our predictions for the NCI/NIH/GSK high confidence collection by only about 2 folds in the recall of biologically similar compounds at a precision of 50% (**Fig. 5d-e**). This relatively poor improvement (compared to that of the RIKEN high confidence collection) was explained by the higher functional diversity of our NCI/NIH/GSK high confidence collection (score of ∼25.3, against ∼14.6 for the RIKEN high confidence collection) although the two collections exhibited similar structural diversity (score of ∼62) (see **Methods**). This high functional diversity of the NCI/NIH/GSK high confidence collection was due to the presence of 6 functionally different sub-collections, which consequently affected the ability of our models to learn chemical-genetic similarities at a high performance for this collection. Although model performance was disturbed by the higher diversity of our NCI/NIH/GSK high confidence collection, we still measured more distinct learning curves from the baseline while labeling 20% of functionally most similar compound pairs as true positives (**Fig. 5e**), indicating that functionally similar pairs were eventually pushed up to the top of the ranked lists by our learning models. Furthermore, we combined the two collections, which added not only more diversity but also more compounds to the RIKEN high confidence collection, and made predictions for the combined dataset, resulting in about a 4.5-fold improvement in the recall of biologically similar compounds at the precision of 50% (**Fig. 5c**). To accomplish higher prediction performance, we, therefore, would need a larger training set (compounds with known chemical-genetic interaction profiles) to compensate for the high functional diversity of compound collections and facilitate the learning process.

**Figure 5.**
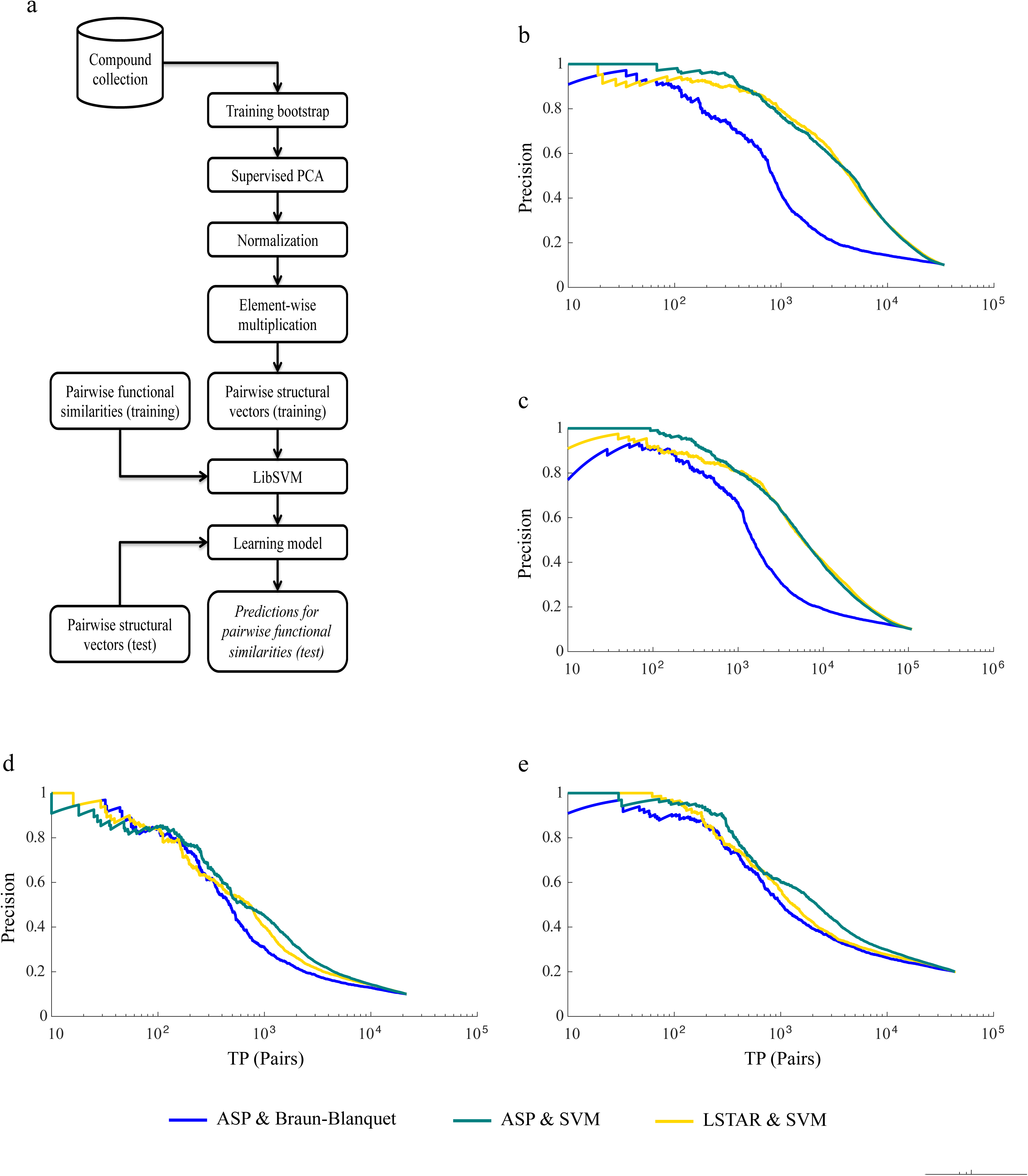
Prediction performance of learning models. **(a)**Learning pipeline for one bootstrap using pairwise structural encodings (see **Methods**). **(b)** Model performance for our RIKEN high confidence collection. The blue curve was the prediction performance of ASP fingerprints paired with the Braun-Blanquet similarity coefficient, whereas the teal and gold curves represented the performance of ASP and LSTAR fingerprints using our learning models, respectively. **(c)** Model performance for the combined RIKEN and NCI/NIH/GSK high confidence collections. **(d)** Performance of the learning models for the NCI/NIH/GSK high confidence collection, where true positives were only 10% (default for this paper) or **(e)** 20% of the compound pairs with the highest chemical-genetic profile similarity.

### Predictive power of structural similarity and SVM models

To investigate the compounds driving our prediction models and the underlying function, we clustered our compound collections into 10 functional as well as 10 structural clusters using K-means and K-medoids, respectively, and mapped only the true positives at the top of our PR curves to their corresponding functional and structural clusters (**Fig. 6**). We found that a large group of compounds generating high prediction scores at the top of our learning curves belonged to the same functional clusters (**Fig. 6b**), whereas the baseline curve for the ASP fingerprints and Braun-Blanquet similarity coefficient included several functional clusters even at lower recalls (**Fig. 6c**). Therefore, our learning models placed more emphasis on the learning of a few certain functional clusters and boosted our prediction performance for those clusters. The first functional cluster that appeared on the learning curve of ASP fingerprints for the RIKEN high confidence collection (blue bar in **Fig. 6b**) was enriched for cell cycle processes based upon the predictions at the MOSAIC database [29] (**Suppl. Tables 3-4** for enrichment of functional clusters). Despite this functional tendency that the learning models showed, several structural clusters contributed to the predicted pairs with high scores for the learning models (**Fig. 6e**), which could lead us to discovering structurally diverse compounds that would exhibit similar biological activities. The discovery of such compounds was of crucial importance since exploring functional analogs with dissimilar structures was entirely out of the capacity of the similar property principle. Hence, our learning models were able to extract compounds with similar function but distinct structures for a target drug/compound, which was far beyond the scope of structural similarity coefficients. For instance, our learning model for the RIKEN high confidence collection assigned a high score to the biologically similar compounds NPD2186 and NPD3120 (chemical-genetic profile similarity of 0.862), while the Braun-Blanquet similarity coefficient for this pair was as low as 0.027 (**Fig. 8c; Suppl. Table 5**). This property of our learning models was achieved by design, where we extracted only a subset of the supervised principal components that were significantly related to the biological activity of compounds, and we then weighted this subset of supervised principal components using support vector regression models. This model property existed in our learning models for the NCI/NIH/GSK high confidence collection as well but in a weaker manner due to the high functional diversity of this collection (**Fig. 7** and **Fig. 8d**).

**Figure 6.**
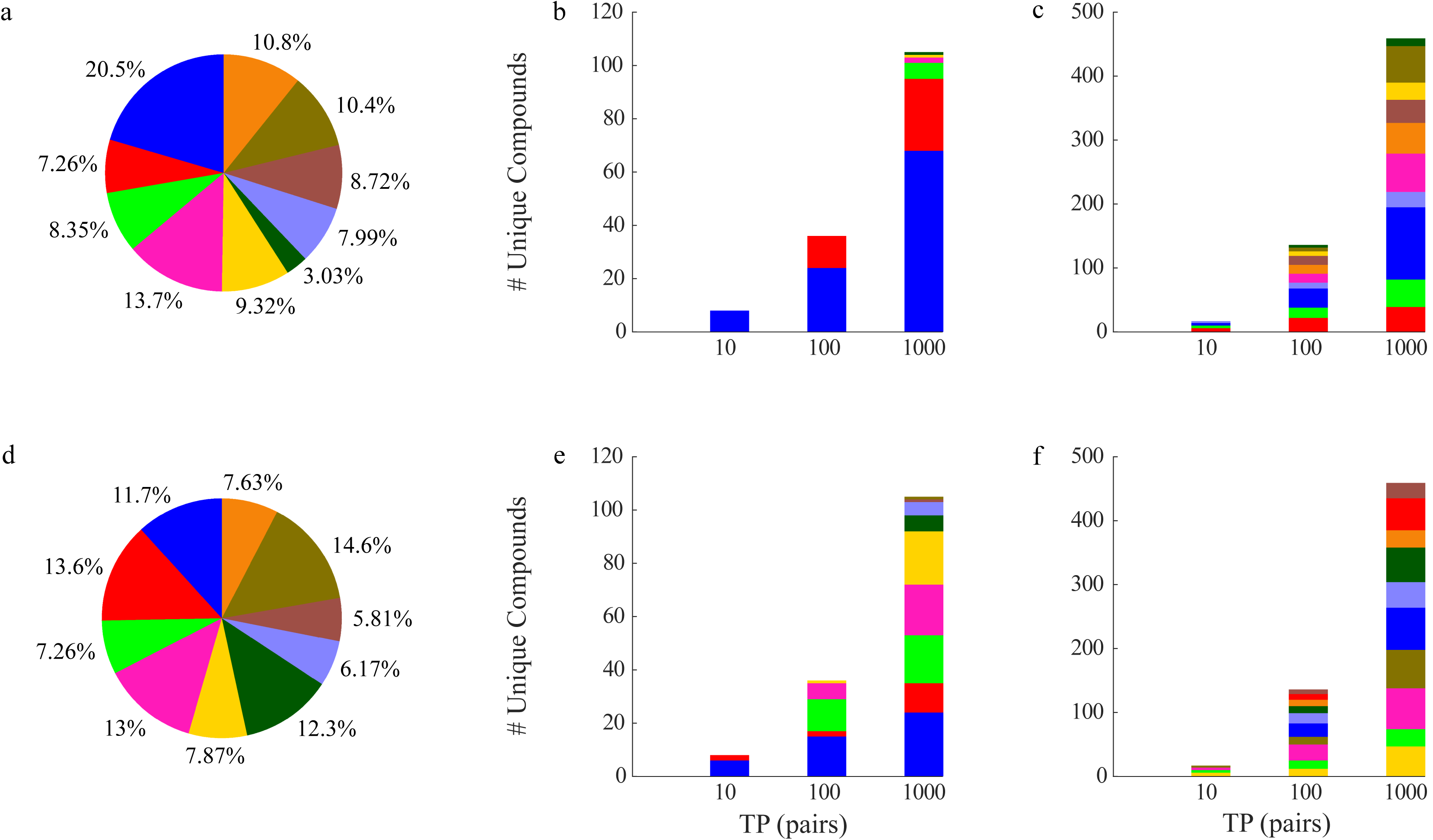
Functional and structural clustering for our RIKEN high confidence collection. **(a)** Distribution of 10 functional clusters generated by k-means using chemical-genetic profiles. Contribution of these functional clusters to the top TP pairs extracted by **(b)** our learning model and **(c)** the Braun-Blanquet similarity coefficient using the ASP fingerprints. **(d)** Distribution of 10 structural clusters generated by k-medoids using the ASP fingerprints. Contribution of these structural clusters to the top TP pairs that were introduced by **(e)** our learning model and **(f)** the Braun-Blanquet similarity coefficient.

**Figure 7.**
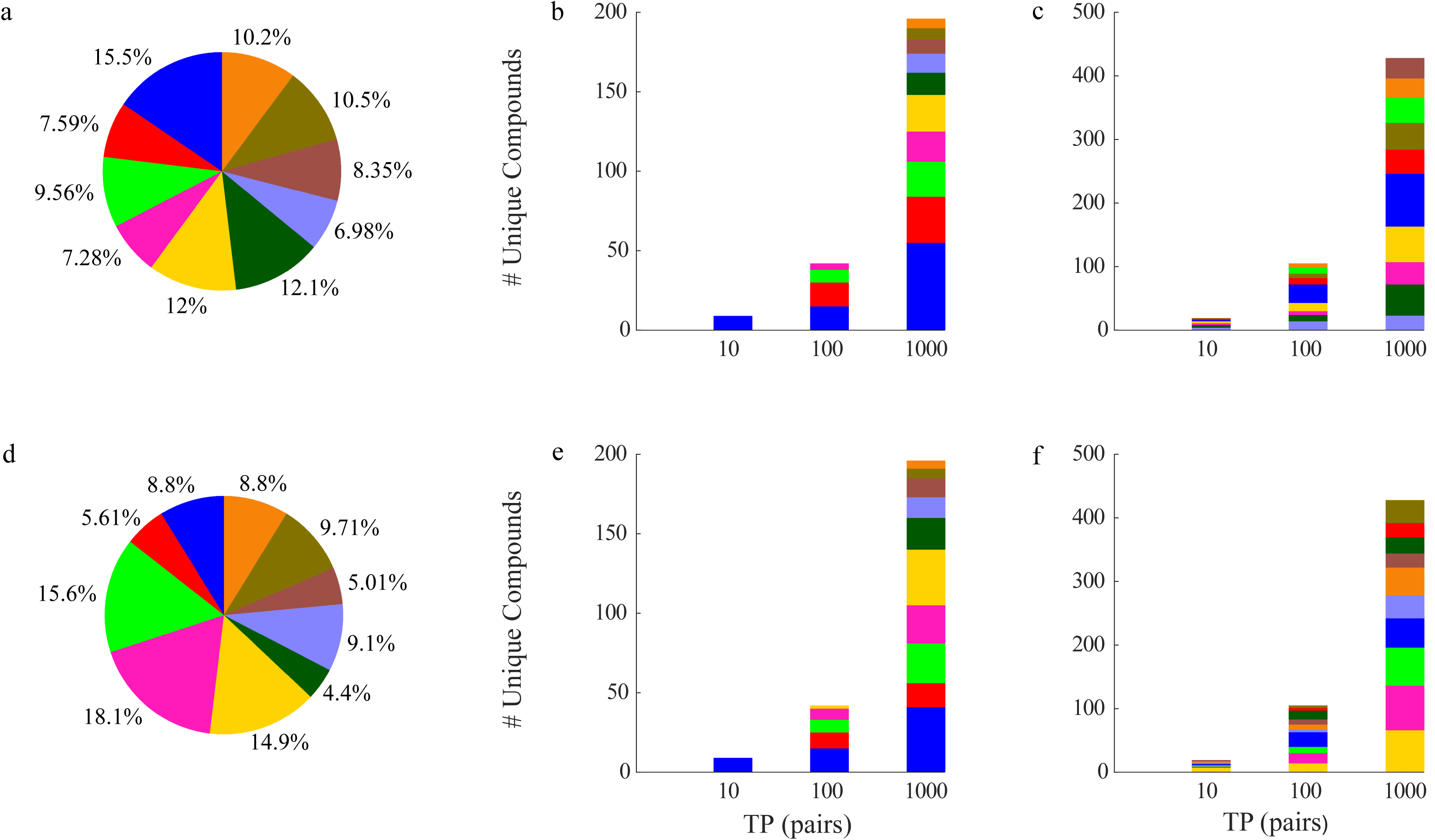
Functional and structural clustering for our NCI/NIH/GSK high confidence collection. **(a)** Distribution of 10 functional clusters generated by k-means using chemical-genetic profiles. Contribution of these functional clusters to the top TP pairs extracted by **(b)** our learning model and **(c)** the Braun-Blanquet similarity coefficient using the ASP fingerprints. **(d)** Distribution of 10 structural clusters generated by k-medoids using the ASP fingerprints. Contribution of these structural clusters to the top TP pairs that were introduced by **(e)** our learning model and **(f)** the Braun-Blanquet similarity coefficient.

**Figure 8.**
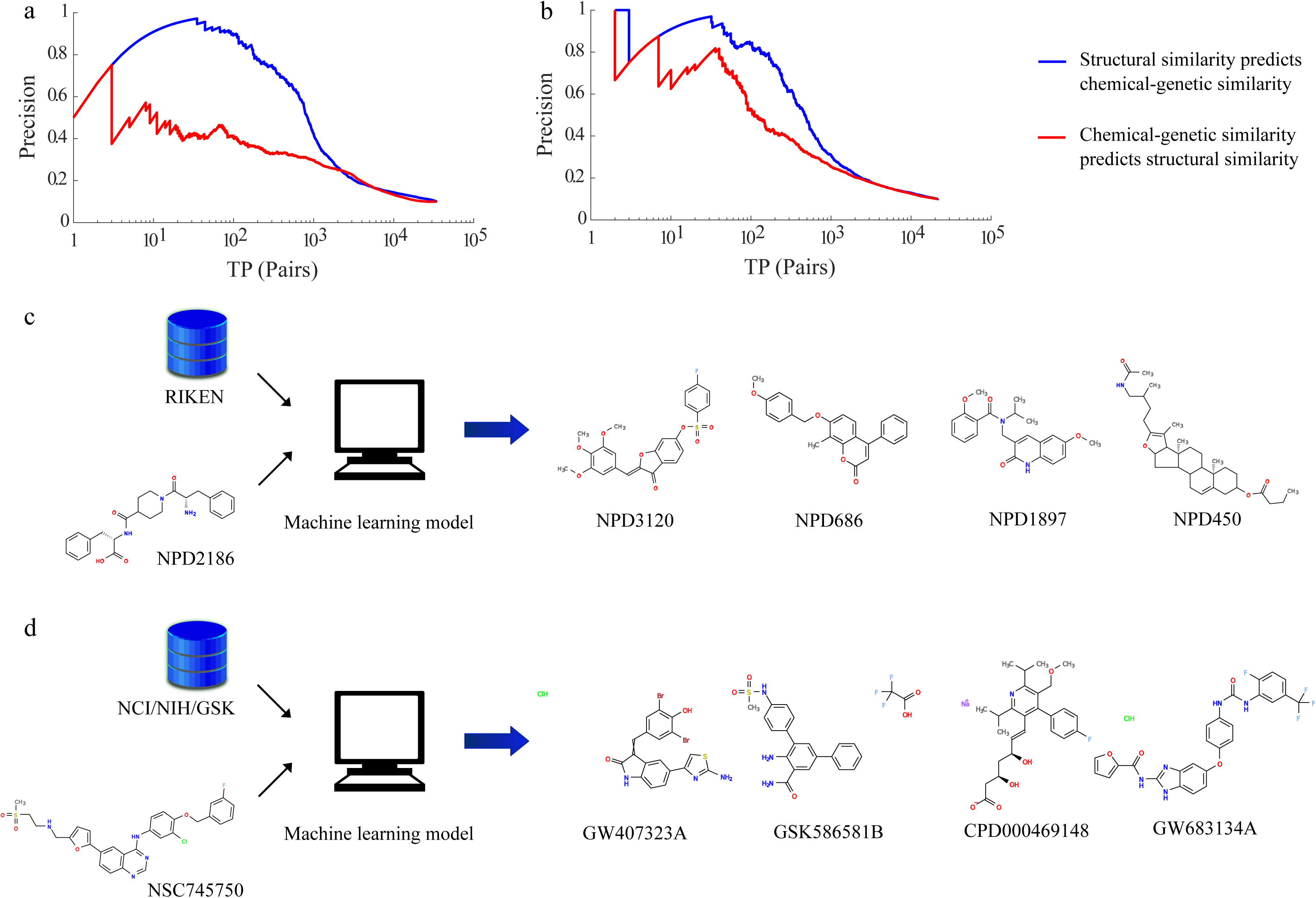
Predictive power of structural similarity as a result of chemical substructures. By using the **(a)** RIKEN and **(b)** NCI/NIH/GSK high confidence collections, we measured that structural similarity showed higher predictive power of chemical-genetic profile similarity than the latter of the former largely because of substructures. Our learning models extracted biologically similar but structurally very dissimilar compounds for **(c)** NPD2186 from our RIKEN high confidence collection and for **(d)** NSC745750 from our NCI/NIH/GSK high confidence collection.

Furthermore, we assessed the predictive power of structural similarity (using the Braun-Blanquet coefficient) against that of chemical-genetic profile similarity for our collections. For the former, we predicted the chemical-genetic profile similarity of our compounds from chemical structures, whereas, for the latter, we used the chemical-genetic profile similarity of compounds to predict their structural similarity. The PR curves revealed that the structural similarity of our compounds had higher predictive power of the chemical-genetic similarity than the latter of the former (**Fig. 8a-b**) since compounds with similar biological activity could represent completely different chemical structures. On the other hand, compounds with similar structures were highly expected to exhibit similar biological activity; therefore, our results were a strong confirmation for the similar property principle. Moreover, the degree of superiority of the predictive power of structural similarity over functional similarity was an indicator of the amount of substructures (i.e., compounds with similar biological activity but distinct structures) that existed in the collection. The wide gap between the curves for the RIKEN high confidence collection (**Fig. 8a**), therefore, represented a large number of substructures in this collection (see **Fig. 6**), while the narrow gap between the curves for the NCI/NIH/GSK high confidence collection (**Fig. 8b**) served as a signal that this collection, which was composed of several sub-collections, included more of one-to-one correspondence between structural and functional profiles. Since the high power of our learning models was to discover compounds of various structures (in addition to compounds with similar structures) that exhibited similar biological activity to a target drug/compound, our learning method showed enormous superiority over the baseline approach (ASP fingerprints paired with the Braun-Blanquet similarity coefficient) for our RIKEN high confidence collection. Although our learning method improved predictions for the functionally diverse collections (such as our NCI/NIH/GSK high confidence collection) moderately, this method exhibited strong predictions for the larger collections representing certain biological functions with structurally diverse compounds.

## CONCLUSION

The chemical informatics community has adopted a broad range of structure descriptors and similarity coefficients for ligand-based virtual screening where the similar property principle has been the basis for ranking of compounds with similar biological activity to a target molecule from chemical structures. However, the research community has lacked a systematic, unbiased benchmark for biological activity that would cover a wide range of targets to definitively assess the performance of alternatives. We generated chemical-genetic interaction profiles from yeast in our labs, covering 13,431 compounds from the RIKEN NPDepo and several NCI/NIH/GSK compound collections, and used these profiles as an unbiased standard for the biological activity of our compounds. Using these chemical-genetic interaction profiles as the basis for the function of our compounds, we systematically benchmarked 10 different structure descriptors and 12 different similarity coefficients. We found that the ASP (and LSTAR) fingerprints paired with the Braun-Blanquet similarity coefficient revealed as the superior choice for ranking of compounds with similar biological activity to a target molecule. The ASP fingerprints encoded all shortest paths between atoms obtained through an exhaustive depth-first search of the molecular graph (up to a predefined depth), and the Braun-Blanquet coefficient represented the number of features shared between two molecules divided by the number of features presented in the larger one. Moreover, we devised a machine learning model that boosted the predictive power of several fingerprints, although the degree of improvement was correlated to the functional diversity of our compound collections. We found that structural similarity had a higher predictive power in prediction of functional similarity than the latter of the former because several substructures contributed to the similar biological activity. Although similarity coefficients predicted the compounds that had both similar function and similar structure, our learning models assigned higher predictive scores to most compounds with similar function by weighting the supervised principal components that were strongly correlated to the chemical-genetic profiles. Therefore, our learning models were able to predict compounds from a library with similar biological activity but diverse structures to a target molecule, which significantly improved performance relative to simple similarity coefficients applied to structure descriptors.

## ACKNOWLEDGEMENTS

We thank Jeff Piotrowski, Sheena Li, and Charlie Boone for constructive feedback on our analysis and results. This work was partially supported by the National Institutes of Health (R01HG005084, R01GM104975) and the National Science Foundation (DBI 0953881). CM is a fellow in the Canadian Institute for Advanced Research (CIFAR) Genetic Networks Program. Computing resources and data storage services were partially provided by the Minnesota Supercomputing Institute and the UMN Office of Information Technology, respectively. SWS was supported by an NSF Graduate Research Fellowship (00039202), an NIH Biotechnology training grant (T32GM008347), and a BICB one-year fellowship.

## MATERIAL AND METHODS

### Data Collections

We used two different compound collections independenty: Our RIKEN high confidence collection, as a subset of the RIKEN NPDepo, was composed largely of purified natural products or natural product derivatives, whereas our NCI/NIH/GSK high confidence collection was a diverse set of several sub-collections: 4 collections from the National Cancer Institute’s Open Chemical Repository (natural products, approved oncology drugs, and structural and mechanistic diversity sets), a library of compounds from the National Institutes of Health Small Molecule Repository with a history of use in human clinical trials (NIH Clinical Collection), and the Glaxo-Smith-Kline kinase inhibitor collection (GSK).

### Designing the support vector machine learning pipeline

We proposed support vector regression (SVR) models for the prediction of the functional activity of our compounds based upon their chemical structures. We used LibSVM [28] for the implementation of our models and bootstrapping [27] for generating our training and test sets. To generate these training and test sets, we drew N (the total number of compounds in a collection) samples uniformly random from the collection but with replacement, assigning ~0.632N unique compounds to the training and the rest to the test set. We used supervised principal component analysis [26] with adaption of chemical-genetic interaction profiles, assuming that these profiles were known for the training set but unknown for the test set, to lower the dimension of structure spaces. We normalized each structural vector that described a compound in the low-dimensional space by its Euclidean length and multiplied each pair of the normalized vectors (both from the training set or both from the test set) in the element-wise manner to create a new structure space, called “pairwise structural encodings”, for the representation of compound pairs (**Suppl. Fig.2a**). We devised a pipeline (**Fig. 5a**) to predict our chemical-genetic profile similarities by using pairwise structural encodings, feeding our regression models with 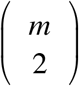 pairwise structural vectors and chemical-genetic profile similarities, where m was the number of compounds in the training set, to predict the chemical-genetic profile similarities for the 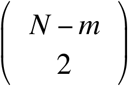 compound pairs that were corresponding to the test set. We used Radial Basis Function (RBF) kernels to build up epsilon support vector regression models and input a number of bootstraps to the models to evaluate the average performance of our models across all bootstraps (**Suppl. Fig. 2b**). To measure the prediction of our pipeline for a newly seen input, we needed to take the average over all the model outputs resulted from different bootstraps, where the new input was treated as a test data in the test sets; the higher the number of bootstraps the more accurate the prediction value. We generated a large number of bootstraps (200 bootstraps for the RIKEN and NCI/NIH/GSK high confidence collections as well as 100 bootstraps for the combined collection) for our evaluations although the performance of our learning models was constant after meeting a certain number of bootstraps.

### Estimating the diversity of compound collections

We assigned all the compounds in a collection to a single cluster and split up the cluster recursively to form clusters of more similar compounds. At any step of recursion, we determined the cluster with the lowest average within-cluster chemical-genetic profile similarity (to compute the functional diversity) or structural similarity (to compute the structural diversity) and divided the cluster into two new clusters using K-means or K-medoids clustering. We stopped generating new clusters right before our algorithm would generate at least two individual clusters exceeding our predefined hard limit for the maximum average between-cluster chemical-genetic similarity (cosine similarity of 0.3) or structural similarity (Braun-Blanquet similarity of 0.3). We repeated the algorithm many times (1000 times for the functional diversity and 100 times for the structural diversity) and computed the mean diversity score as the average exponentiation of the Shannon entropy indices over all the instances:

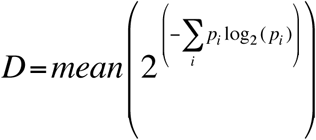

where *p_i_* was the proportional abundance of compounds in the *i^th^* cluster of the final clustering.

---

**Suppl. Figure 1. Depth impact of structure descriptors on the performance of prediction models.** We measured the precision of our prediction models at 10 consequent molecular depths for 5 different structure descriptors, each paired with the Braun-Blanquet similarity coefficient, using our NCI/NIH/GSK high confidence collection.

**Suppl. Figure 2. Pairwise structural encodings and bootstrapping. (a)** Pairwise structural features created by the element-wise multiplication of the normalized, low-dimensional structural vectors. We reduced dimension of descriptors using a supervised principal component analysis method. **(b)** Smoothing average over bootstraps. At each bootstrap, the chemical-genetic profile similarity of the test pairs (represented by “X”) was predicted, and all the predicted values for a test pair at different bootstraps were averaged to smoothen the prediction. For example, the compound pair 1 was a test pair in bootstraps 1, 2, and 4 (In bootstrap 3, it might be a training pair or an invalid pair where one compound belonged to the training set and the other to the test set).

**Suppl. Table 1. Precision at several recalls for all combinations of structure descriptors and similarity coefficients using our RIKEN high confidence collection.**

**Suppl. Table 2. Precision at several recalls for all combinations of structure descriptors and similarity coefficients using our NCI/NIH/GSK high confidence collection.**

**Suppl. Table 3. Functional enrichment of clusters for the 1000 top TP pairs of our learning model using our RIKEN high confidence collection.**

**Suppl. Table 4. Functional enrichment of clusters for the 1000 top TP pairs of our learning model using our NCI/NIH/GSK high confidence collection.**

**Suppl. Table 5. The list of compounds with similar chemical-genetic interaction profiles but dissimilar chemical structures for the largest functional cluster of 1000 top TP pairs (blue bars in Figs. 6b–7b).**

## References

1. Scannell JW, Blanckley A, Boldon H, Warrington B. Diagnosing the decline in pharmaceutical R&D efficiency. Nat. Rev. Drug Discov. 2012; 11: p. 191–200.

2. Johnson MA, Maggiora GM. Concepts and applications of molecular similarity: John Wiley; 1990.

3. Bajorath J. Integration of virtual and high-throughput screening. Nat. Rev. Drug Discov. 2002; 1: p. 882–894.

4. Tanrikulu Y, Krüger B, Proschak E. The holistic integration of virtual screening in drug discovery. Elsevier Drug Discov. Today. 2013; 18: p. 358–364.

5. Stahura FL, Bajorath J. Virtual Screening Methods that Complement HTS. Combinatorial Chemistry & High Throughput Screening. 2004; 7: p. 259–269.

6. Eckert H, Bajorath J. Molecular similarity analysis in virtual screening: foundations, limitations, and novel approaches. Elsevier Drug Discov. Today. 2007; 12: p. 225–233.

7. Guha R, Van Drie JH. Structure-Activity Landscape Index: Identifying and Quantifying Activity Cliffs. J. Chem. Inf. Model. 2008; 48: p. 646–658.

8. Bajorath J, Peltason L, Wawer M, Guha R, Lajiness MS, Van Drie JH. Navigating structure–activity landscapes. Elsevier Drug Discov. Today. 2009; 14: p. 698–705.

9. Wassermann AM, Wawer M, Bajorath J. Activity Landscape Representations for Structure-Activity Relationship Analysis. J. Med. Chem. 2010; 53: p. 8209–8223.

10. Duanb J, Dixona SL, Lowriea JF, W. S. Analysis and comparison of 2D fingerprints: Insights into database screening performance using eight fingerprint methods. J. Molecular Graphics and Modelling. 2010; 29: p. 157–170.

11. Willett P. Similarity-based virtual screening using 2D fingerprints. Elsevier Drug Discov. Today. 2006; 11: p. 1046–1053.

12. Raymond JW, Willett P. Effectiveness of graph-based and fingerprint-based similarity measures for virtual screening of 2D chemical structure databases. J. Computer-Aided Molecular Design. 2002; 16: p. 59–71.

13. Willett P. Similarity searching using 2D structural fingerprints. In.: Springer; 2011. p. 133–158.

14. Fligner MA, Verducci JS, Blower PE. A Modification of the Jaccard–Tanimoto Similarity Index for Diverse Selection of Chemical Compounds Using Binary Strings. Technometrics. 2002; 44: p. 110–119.

15. Holliday JD, Salim N, Whittle M, Willett P. Analysis and Display of the Size Dependence of Chemical Similarity Coefficients. J. Chem. Inf. Comput. Sci. 2003; 43: p. 819–828.

16. Giaever G, et al. Functional profiling of the Saccharomyces cerevisiae genome. Nature. 2002; 418: p. 387–391.

17. Parsons A, et al. Exploring the mode-of-action of bioactive compounds by chemical-genetic profiling in yeast. Cell. 2006; 126: p. 611–625.

18. Giaever G, et al. Chemogenomic profiling: Identifying the functional interactions of small molecules in yeast. Proc. Natl. Acad. Sci. U. S. A. 2004; 101: p. 793–798.

19. Parsons A, et al. Integration of chemical-genetic and genetic interaction data links bioactive compounds to cellular target pathways. Nat. Biotech. 2004; 22: p. 62–69.

20. Piotrowski JS, et al. Functional annotation of chemical libraries across diverse biological processes. Nat. Chem. Bio. (under review).

21. Vapnik VN. Statistical Learning Theory: John Wiley; 1998.

22. Simpkins SW, et al. in preparation.

23. Hinselmann G, Rosenbaum L, Jahn A, Fechner N, Zell A. jCompoundMapper: An open source Java library and command-line tool for chemical fingerprints. J. Cheminformatics. 2011; 3: p. 1–14.

24. Rogers D, Hahn M. Extended-connectivity fingerprints. J. Chem. Inf. Model. 2010; 50: p. 742–754.

25. Hayek LC. Measuring and monitoring biological diversity: standard methods for amphibians. In Heyer WR, et al., editors.: Smithsonian Books, Washington, D.C.; 1994.

26. Barshan E, Ghodsi A, Azimifar Z, Zolghadri Jahromi M. Supervised principal component analysis: Visualization, classification and regression on subspaces and submanifolds. Pattern Recognition. 2011; 44: p. 1357–1371.

27. Efron B, Tibshirani R. Improvements on cross-validation: the 632+ bootstrap method. J. American Statistical Association. 1997; 92: p. 548–560.

28. Chang CC, Lin CJ. LIBSVM: a library for support vector machines. ACM Transactions on Intelligent Systems and Technology (TIST). 2011; 2: p. 1–27.

29. Nelson J, et al. in preparation.

